# Dysfunctional effector memory CD8 T cells in the bronchoalveolar compartment of people living with HIV

**DOI:** 10.1101/2023.05.05.539571

**Authors:** Maphe Mthembu, Helgard Claassen, Sharon Khuzwayo, Valentin Voillet, Anneta Naidoo, Kennedy Nyamande, Dilshaad F. Khan, Priya Maharaj, Mohammed Mitha, Zoey Mhlane, Farina Karim, Erica Andersen-Nissen, Thumbi Ndung’u, Gabriele Pollara, Emily B. Wong

**Author notes:** Correspondence: Dr Emily Wong, Africa Health Research Institute, Durban, South Africa and Dr Gabriele Pollara, University London College, London, UK. contributed equally to this work (Gabriele Pollara, Emily B. Wong).

## Abstract

Mechanisms by which HIV causes susceptibility to respiratory pathogens remain incompletely understood. We obtained whole blood and bronchoalveolar lavage (BAL) from people with latent TB infection in the presence or absence of antiretroviral-naïve HIV co-infection. Transcriptomic and flow cytometric analyses demonstrated HIV-associated cell proliferation plus type I interferon activity in blood and effector memory CD8 T-cells in BAL. Both compartments displayed reduced induction of CD8 T-cell-derived IL-17A in people with HIV, associated with elevated T-cell regulatory molecule expression. The data suggest that dysfunctional CD8 T-cell responses in uncontrolled HIV contribute to susceptibility to secondary bacterial infections, including tuberculosis.

## Introduction

People living with HIV (PLWH), have >18-fold increased risk of respiratory infections, including tuberculosis (TB) (1). Uncontrolled HIV infection results in several immune perturbations, including CD4 T-cell depletion, cytotoxic CD8 T-cell expansion (2) and elevated cytokine activity (3) (e.g. type I interferons (4)). Most studies have focused on immune perturbations in blood but less on target organs for co-infections, such as the lungs. Improved understanding of HIV’s impact at the site of host-pathogen interactions may offer insights into immune modulation caused by uncontrolled HIV infection, and in turn, identifying putative therapeutic targets in PLWH. Whole compartment transcriptional profiling permits unbiased assessments of disease-mediated perturbation by HIV at the molecular level, whilst still retaining the ability to deconvolute differences in immune cell frequency, cytokine activity and other functions (5, 6).

In this study, we aimed to explore the lung mucosal immune landscape, through transcriptomic profiling of bronchoalveolar lavage (BAL) cells in individuals with untreated HIV infection. We previously established the feasibility of this technique in a small cohort (7). The current study extends this approach in a population with evidence of *Mycobacterium Tuberculosis* (*Mtb*) infection (i.e. latent TB) in a high TB transmission setting. We test the hypothesis that HIV results in compartment-specific immune perturbations that may underlie the susceptibility to TB disease and other respiratory infections.

## Methods

### Study subjects and samples collection

We recruited otherwise healthy individuals grouped by HIV serostatus, from a previously described research bronchoscopy cohort in Durban, South Africa (7). Inclusion criteria included a positive QuantiFERON-TB Gold Plus for all participants and, for HIV seropositive individuals, no previous exposure to antiretroviral therapy (ART). Paired blood and BAL samples were obtained from all study participants, and peripheral blood mononuclear cells (PBMCs) and BAL mononuclear cells (BLCs) isolated (Supplementary Methods).

### Flow cytometry

Mononuclear cells from both compartments were stained with monoclonal antibodies and some were stimulated with a non-specific mitogen and further stained for intracellular cytokines. All cells were then acquired, and some subsets sorted through BD FACS™ Aria III (detailed in Supplementary Methods).

### Transcriptional profiling

RNA was extracted from cryopreserved whole compartment BAL and blood samples, as well as sorted cell populations in BAL. cDNA libraries were prepared and sequenced on an Illumina NextSeq 500 platform (details in Supplementary Methods).

RNA sequences were quantified, aligned, and annotated to measure and deduce patterns of gene expression. Whole-genome differential gene expression was analysed. Transcriptional module expression was quantified as previously described (5), and differential module enrichment between groups was assessed using Mann-Whitney test (Supplementary Methods).

## Results

Thirty-seven participants contributed paired samples, 70% female, with median age 32 (IQR 25-35, Supplementary Table 1). Sex and age distribution were comparable between PLWH and those without HIV. Median CD4 count in PLWH was 513 cells/mm^3^ (IQR 275-681) and median VL 25,229 copies/mL (IQR 13,987-33,565).

Unsupervised gene expression analyses identified 139 upregulated and 44 downregulated genes in blood of PLWH (Figure 1A, left), in contrast to 13 upregulated and 1 downregulated, in BAL (Figure 1A, right). Of the 152 genes elevated in PLWH across both compartments, only 3 (2%) were increased in both blood and BAL (Figure 1B), highlighting the compartmentalized nature of HIV-induced transcriptional changes. HIV-associated changes in blood mapped to cell proliferation, interferon signaling and antigen presentation pathways, whereas in BAL, immunoregulatory pathways were enriched (Figure 1C). Cytokines predicted to regulate upregulated genes in HIV also demonstrated compartment-specific effects, with only 3/12 cytokines (IL-2, IL-15 and IL-12B) upstream of transcripts elevated by HIV in both compartments (Figure 1D).

**Figure 1:**
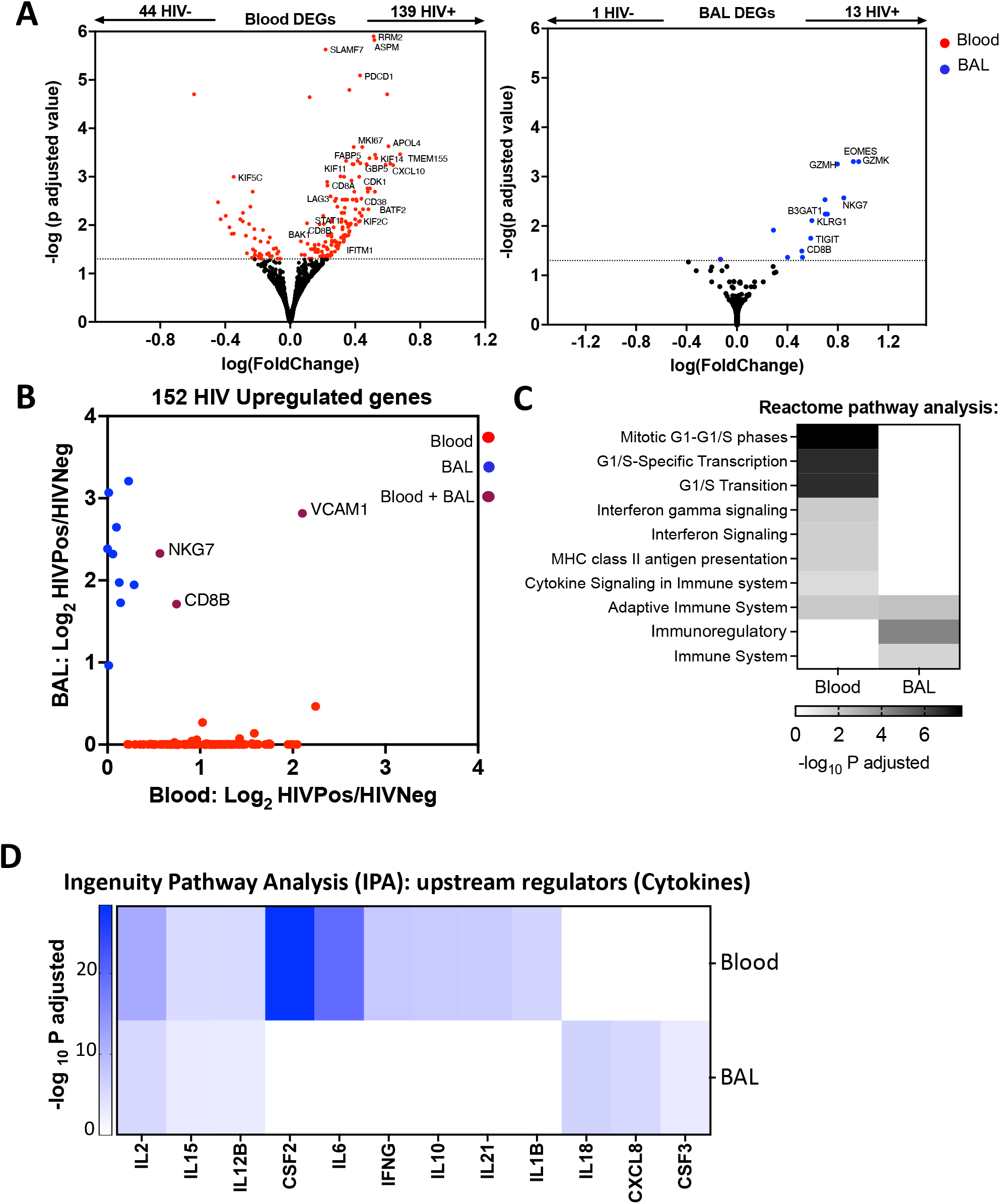
HIV infection associated with different transcriptional profiles in blood compared to the bronchoalveolar space. **A**. Whole genome differential gene expression transcriptional analysis from 10 pairs of blood and bronchoalveolar lavage (BAL) samples from participants with and without HIV (n = 20, HIV-negative (10) and HIV-positive (10)); volcano plots indicate genes with significantly elevated expression in HIV infection in blood (red) and BAL (blue) compartments. **B**. Dot plot displaying adjusted P values of genes with significantly elevated expression in HIV-negative compared to HIV-positive in blood (red), BAL (blue) and both (magenta) compartments. **C**. Heatmap showing REACTOME levels (white: lowest to black: highest) of enriched pathways in PLWH transcriptome blood (left column) and BAL (right column) samples. **D**. Heatmap displaying levels (white: lowest to blue: highest) of upstream cytokines significantly predicted to regulate differential gene expression in PLWH blood and BAL samples.

BAL was predominantly composed of alveolar macrophages (Supplementary Figure 1), but several of the 13 genes overexpressed in PLWH were lymphocyte-associated (Figure 1A, right). Validated transcriptional modules that quantify the relative abundance of immune cells demonstrated relative depletion of CD4 T-cell transcripts in both blood and BAL in PLWH (5) (Figure 2A, left), an expansion of CD8 T-cells in both compartments and no differences in NK cell frequency (Figure 2A, left). These modules were derived from peripheral blood cells, and may not accurately reflect the transcriptional state of BAL lymphocytes (5). Therefore, we generated an independent set of modules derived from genes differentially expressed in sorted BAL cells (gating strategy in Supplementary Figure 2). The BAL-derived module specificity was comparable to those from blood, with better discrimination of CD8 T-cells from MAIT cells (Supplementary Figure 3). These BAL-derived modules confirmed enrichment of CD8 T-cells, but not MAIT cells, in PLWH (Figure 2A, right). Flow cytometry confirmed elevated CD8 T-cell frequency in HIV (Supplementary Figure 4), in turn validating our approach to use transcriptional modules to quantify cell enrichment from BAL and blood transcriptomes.

**Figure 2:**
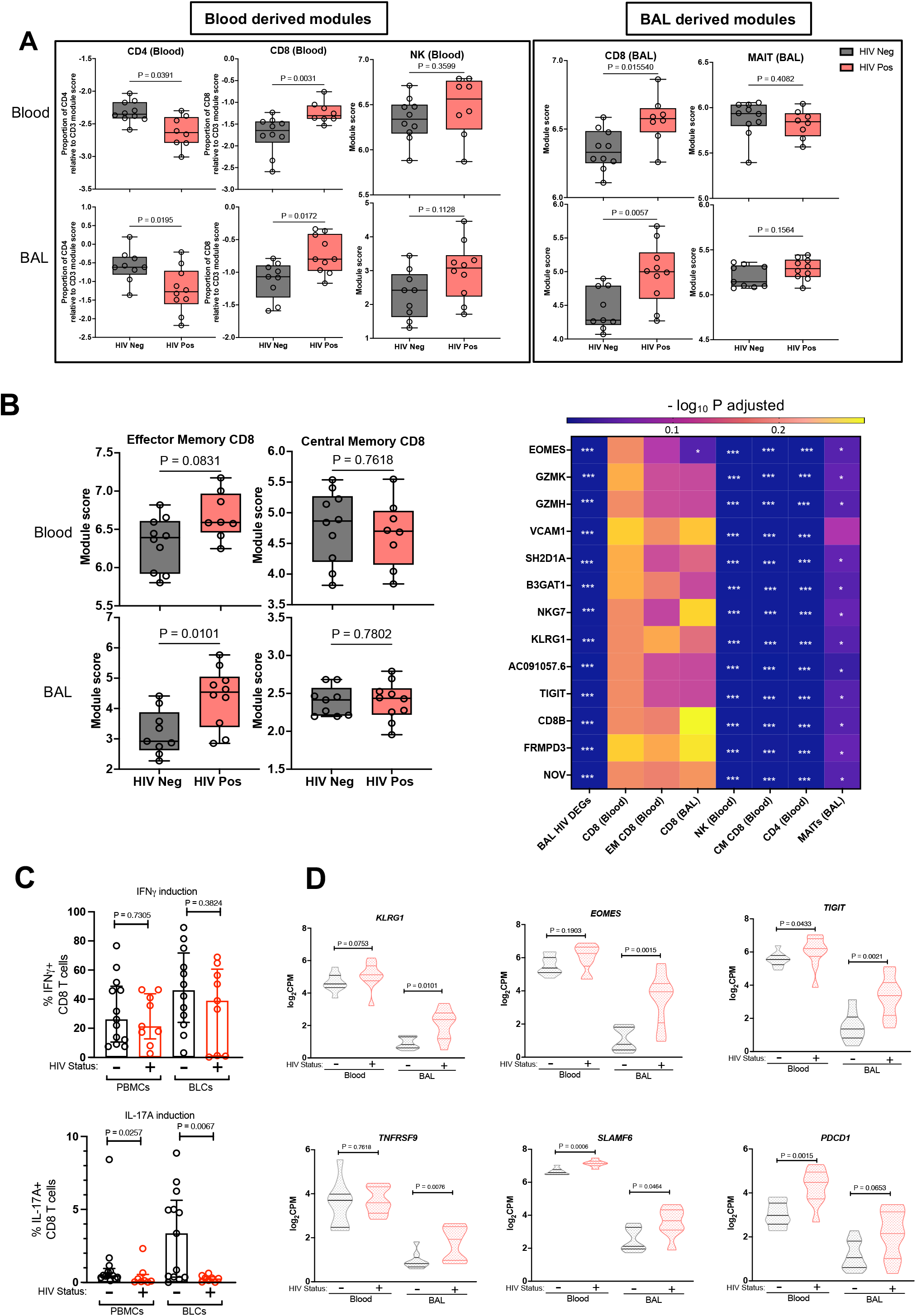
Bronchoalveolar compartment in HIV infection characterized by enrichment of effector memory CD8 T-cells deficient in IL-17A production. Transcriptional module analysis of the whole transcriptomics data using **A**. Blood and BAL-derived modules reflecting immune cell relative frequency, and **B**. effector and central memory CD8 T-cell modules **(left column)** and a heatmap showing adjusted p-values (blue = lowest, yellow = highest, and asterisks representing the level of significance) for 13 genes upregulated in the BAL between PLWH and those without HIV (first column) regressed against 7 transcriptional modules: CD8 T-cells, effector memory CD8 T-cells (EM), central memory CD8 T-cells (CM), CD4 T-cells, NK cells and MAITs cells **(right column). C**. Expression of IFN-γ and IL-17A following PMA-Ionomycin stimulation of mononuclear cells from both blood and BAL (top row). **D**. Immune checkpoint molecule expression in blood and BAL (middle and bottom rows) (n=20, 10 HIV negative and 10 HIV positive).

Several of the 13 genes with higher expression in BAL of PLWH are associated with CD8 T-cell effector functions (*GZMK, GZMH, B3GAT1, NKG7*) and we sought to test the hypothesis that this increased expression reflects an increased frequency of effector memory (EM) CD8 T-cells. We generated and validated new transcriptional modules from different CD8 T-cell functional states (Supplementary Figure 5) and they showed enrichment of CD8 T-cells with an EM phenotype in BAL of PLWH, with no differences in central memory (CM) CD8 T-cells (Figure 2B, left column). To determine whether EM CD8 T-cells were the primary source of significantly upregulated genes in BAL of PLWH, we sought to adjust for differences in cell frequency by performing a linear regression at the whole transcriptome level, controlling for cellular module expression in each sample. This revealed that differential expression of the 13 genes elevated in BAL of PLWH was no longer present following regression against either blood/BAL CD8 T-cell modules or the CD8 EM module, but this was not the case with CD8 CM, CD4, NK or MAIT cell modules (Figure 2B, right column). We interpreted these data to suggest transcriptomic perturbations in untreated PLWH in BAL are likely attributable to increased numbers of EM CD8 T-cells.

Finally, we sought to determine whether the differential frequency of CD8 T-cells in unstimulated BAL in HIV was also associated with different functional activity of these cells in HIV. We quantified production of IFN-γ and IL-17A by both blood and BAL CD8 T-cells before and after stimulation with PMA/ionomycin (Supplementary Figure 6A). Baseline cytokine expression was low and did not differ by HIV status (Supplementary Figure 6B), but following stimulation, we observed a decreased frequency of CD8 T-cells expressing IL-17A in both blood and BAL compartments of PLWH, whereas no such difference was observed for IFN-γ (Figure 2C). Notably, 3/13 of differentially upregulated genes in BAL of PLWH (*KLRG1, EOMES* and *TIGIT*), along with *TNFRSF9* and *SLAMF6*, are associated with CD8 T-cell exhaustion or functional inhibition (8) (Figure 2D). In blood of PLWH, we observed more CD8 T-cell exhaustion markers, *PDCD1* and additionally *TIGIT, SLAMF6*, also showing increased expression indicating a compartment-specific effect (Figure 2D). Flow cytometry analysis confirmed a significant increase in the expression of PD-1 (encoded by the gene *PDCD1*) on blood CD8 T-cells in PLWH. A higher expression of PD-1 on CD8 T-cells in PLWH was seen in BAL but this was not statistically significant (Supplementary Figure 6C). Overall, these results suggest increased immunoregulatory activity in HIV infection that may attenuate IL-17A responses of CD8 T-cells in both blood and BAL.

## Discussion

HIV infection increases the risk of respiratory infections including TB (1). Of the mechanisms proposed to date, most have not included systems-level analysis of the immunobiology in lungs. We demonstrate that uncontrolled HIV infection results in transcriptional changes in the lungs that differ from those in blood. The BAL of PLWH is enriched for CD8 T-cells that transcriptionally resemble EM cells but these show deficient IL-17A responses upon stimulation, a process associated with elevated expression of immunoregulatory molecules. Our data support a model whereby impaired CD8 T-cell production of IL-17A may contribute to susceptibility to secondary bacterial infections in HIV infection (9).

We extend our earlier report in which we first reported bulk transcriptomic assessments from BAL, by including a larger number of participants in whom active lung disease (other than HIV) was rigorously ruled out, and all of whom had evidence of past exposure to *Mtb*. As expected, several pathways were upregulated in the peripheral blood compartment of PLWH including cell proliferation, possibly reflecting a homeostatic response to the depletion of CD4 T-cells in the HIV infection (10), and type 1 and II IFN signaling antiviral pathways. In contrast, we observed markedly fewer differentially expressed genes in the BAL compartment of PLWH compared to HIV-negative controls. This may have reflected the predominance of macrophages in BAL, which do not show transcriptomic perturbation in PLWH at baseline and may have reduced the sensitivity to detect effects on other cell types. Nevertheless, the few genes with elevated expression in PLWH were markedly associated with lymphocyte function, which drew us to focus on T-cell perturbations in the BAL compartment.

Using module analysis as an approach to bioinformatically de-convolute whole-genome transcriptomic data, we confirmed the well-known consequences of HIV infection in depleting CD4 T-cells and enriching CD8 T-cells in both blood and BAL (11). The increased frequency of CD8 T-cells in BAL is consistent with reports of lymphatic alveolitis (12) and was associated with elevated expression of several genes linked to cytotoxicity, such as granzymes H and K, and a transcriptional module reflective of EM CD8 T-cells, but not CM CD8 T-cells, NK or MAIT cells. Nevertheless, both blood and BAL CD8 T-cells showed diminished IL-17A production after ex vivo stimulation, but no impairment in IFN-γ production. This IL-17A defect has been reported in the blood of virally suppressed people with HIV (13), but our current study shows that this is also evident in the airways of people with untreated HIV.

IL-17A is reported to confer protection against hypervirulent *Mtb* strains such as HN878 (14) and this may contribute to susceptibility to TB disease in ARV-naïve PLWH (15). Nevertheless, given that active TB is characterized by exaggerated IL-17A responses that may contribute to tissue pathology (6), our observations may also provide an explanation for why severe chest-X-ray abnormalities, such as cavities, are less common in TB patients with advanced HIV (15). Notably, we observed the defect in IL-17A production in PLWH to be associated with increased expression of molecules associated with T-cell immunomodulation, suggesting an exhausted CD8 T-cell phenotype. Indeed, it has been proposed that elevated expression of inhibitory receptors is responsible for diminished IL-17A responses in HIV (13). Our findings suggest that modulation of negative regulators of T-cell function, with precisely targeted biologics, could be one strategy to restore HIV-induced dysregulated immunity in the lung mucosa.

Our study has several limitations. First, this was a single-center study with a limited sample size that assessed individuals at a single timepoint. Participants with HIV were all ART-naïve, and we cannot predict if the immune perturbations observed would persist during suppression of HIV replication by ART. Additionally, assessment of unstimulated transcriptomes at bulk cell level precluded assessment of the dynamic nature of responses in response to bacterial stimulation. Finally, we only investigated a limited number of cytokine responses at the protein level in blood and BAL cells, leaving unanswered whether the observed defects in IL-17A responses are associated with alterations to a larger set of cytokine responses in HIV still able to leverage.

Our study leveraged unique access to both blood and BAL samples of a well-defined cohort of individuals to unmask immune dysregulation associated with uncontrolled HIV infection at the site of pulmonary host-pathogen interactions. We conclude that HIV is associated with divergent transcriptional effects in the blood and in the bronchoalveolar compartments. Blood was characterized by elevated cell proliferation and type I/II interferon signaling activity, while BAL was enriched for CD8 T-cells with an effector phenotype but a defect in inducible IL-17A production, associated with increased expression of T-cell inhibitory molecules. Future work will need to assess the ability of ART to reverse this immune dysregulation and to explore the role of adjunctive immune checkpoint inhibitors to restore mucosal CD8 T-cell function and improve immune protection against respiratory infections in HIV.

## Supporting information

BAL HIVup DEGs

Blood HIVup DEGs

BAL Modules gene list

## Data availability

Supplementary materials are provided (https://figshare.com/s/27bbd4134c2639d5c7ed) and they consist of data provided by the authors to benefit the reader. The original RNA-Seq datasets generated for this study can be found in the Gene Expression Omnibus (GEO) repository, accession number GSE230738 for whole sample transcriptomics and GSE231628 for mini-population transcriptomics. The supplementary materials are not copyedited and are the authors’ sole responsibility, so questions or comments should be addressed to the corresponding authors.

## Acknowledgments

We appreciate the study participants for their participation in the research, all the nurses from Inkosi Albert Luthuli Hospital for their support in acquiring the blood and BAL samples and the AHRI laboratory biorepository department for handling and distributing the samples.

