## Supplementary Material for "Dysfunctional effector memory CD8 T cells in the bronchoalveolar compartment of people living with HIV"

**Supplementary Table 1: Demographic and clinical characteristics of the enrolled participants**

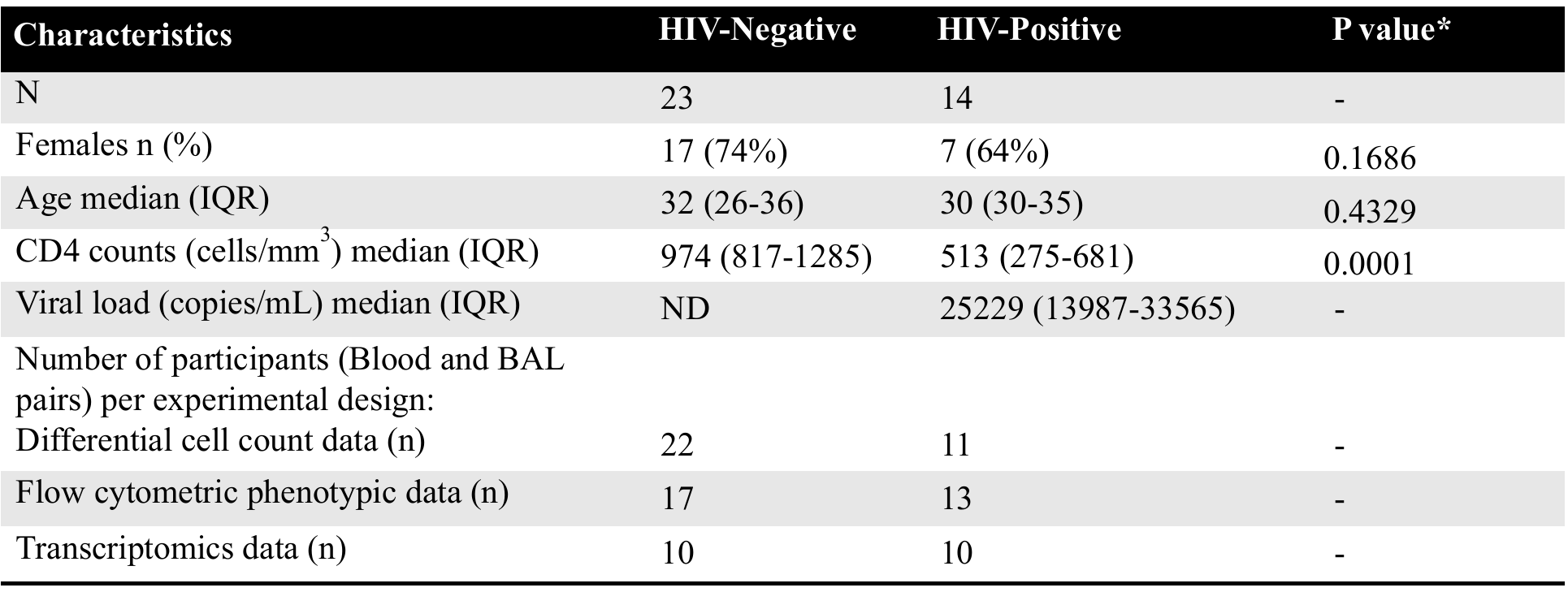

*Mann-Whitney between HIV-negative and HIV-positive people, ND (not detected)

**A	B**

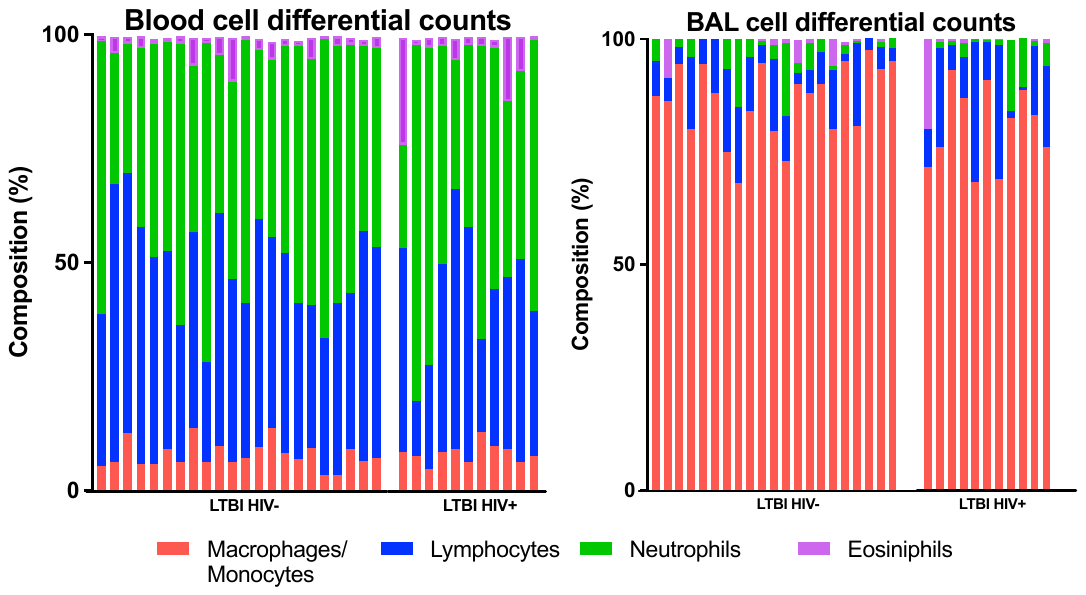

**Supplementary Figure 1: Differential cell counts proportions showing the distribution of the major immune cell types in blood and Bronchoalveolar lavage (BAL) compartment.** **A.** shows proportions in the percentage of the different immune cell major cell types per participant (column) in the blood compartment which are also separated by HIV status on the x-axis. **B.** same proportions of the different immune cells in the BAL compartment.

**A	B	C	D	E	F**

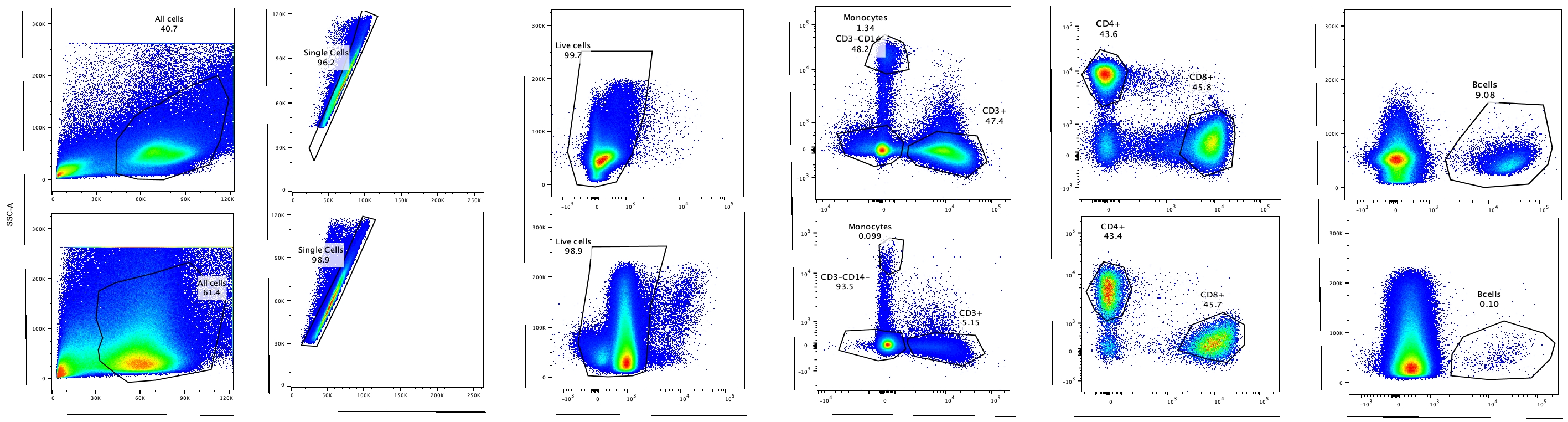

SSC-A

Lymphocytes and Monocytes gate

Dead staining

FSC-A

FSC-H

SSC-A

CD14+

CD3+

CD8+

CD4+

SSC-A

CD19+

Blood

BAL

**Supplementary Figure 2: Flow cytometry gating strategy for confirmation of immune cell subsets. A**. side-scatter-area (SSC-A) vs forward-scatter-Area (FSC-A) are plotted showing the expanded lymphocyte gate which also includes larger-sized monocytes. **B.** FSC-A vs. FSC-Height (-H) are plotted to exclude clustered cells and select for single cells. **C.** dead cell exclusion gate. **D.** CD14+ monocytes, CD3+ T-cells and those that are double negative for CD14 and CD3. **E.** Use cells from the CD3+ T-cells gate to select CD4+ and CD8+ T-cells. **F.** Use double negative cells from the double negative gate and select CD19+ B-cells. This gating strategy shows representative gates from stained mononuclear cells from both blood: top row and BAL: bottom row.

**A**

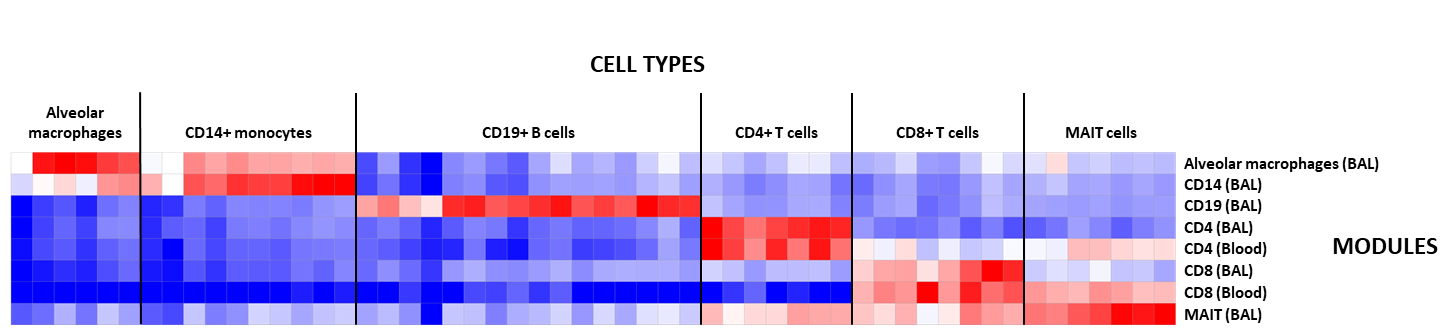

**B**

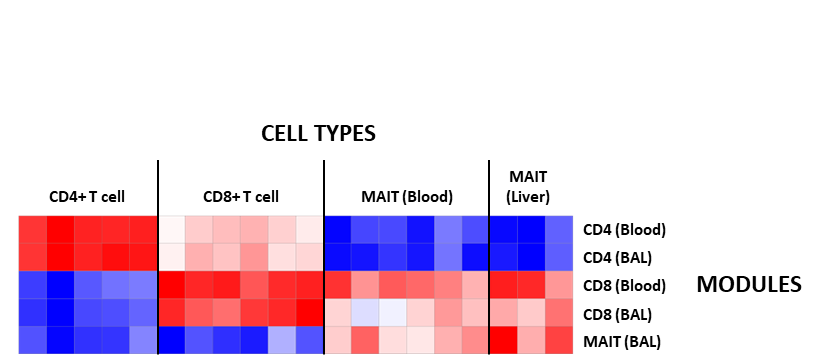

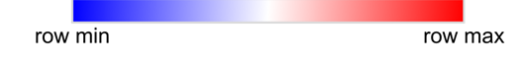

**Supplementary Figure 3: Assessment of sensitivity and specificity of Bronchoalveolar Lavage (BAL)-derived immune cell modules.** The expression of transcriptional modules derived from flow cytometry sorted immune BAL cells in (A) BAL samples in HIV-positive individuals in the current cohort and (B) sorted CD4, CD8 and MAIT cells (ArrayExpress dataset E-MTAB-7143).

**A	B	C**

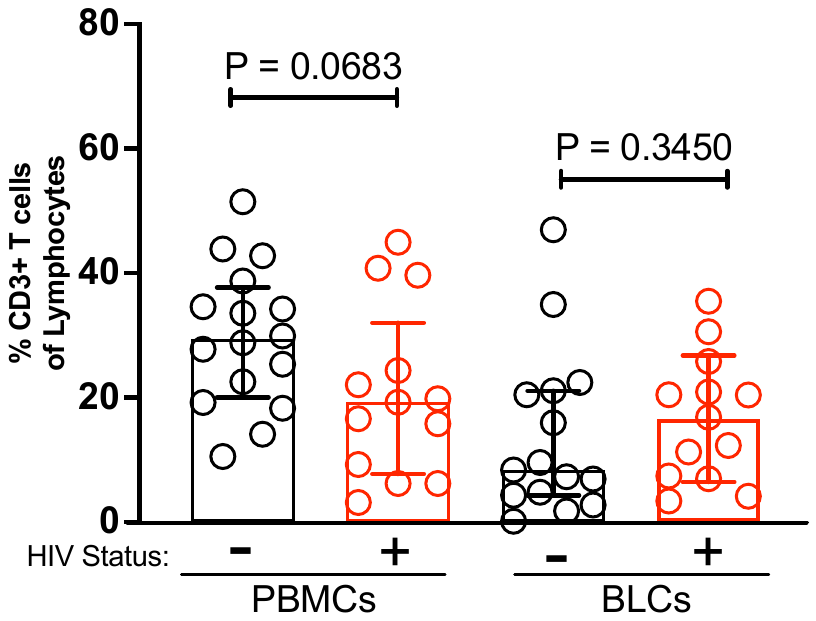

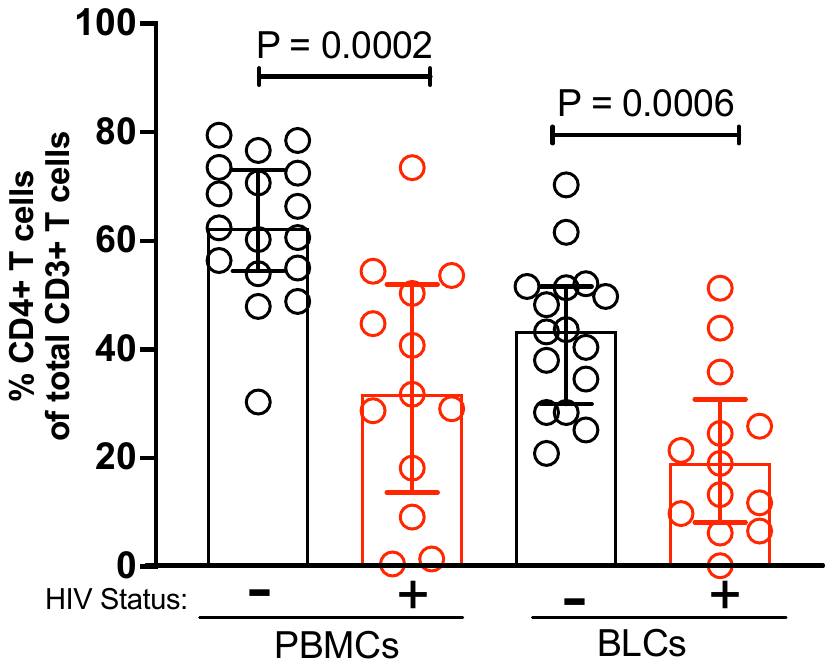

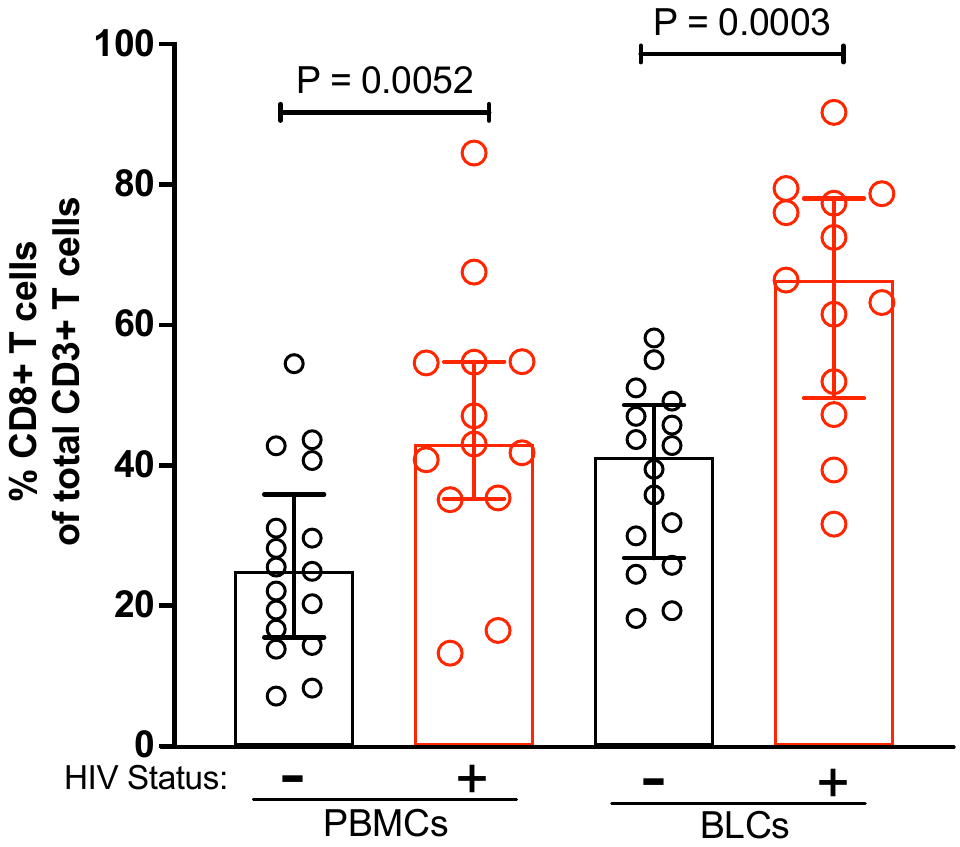

**Supplementary Figure 4: Enumeration of immune cells in Peripheral Blood Mononuclear Cells (PBMCs) and Bronchoalveolar Lavage mononuclear Cells (BLCs) in PLWH and those without HIV.** Quantification of (A) T cells by expression of CD3 T cells, (B) CD4 T cells by expression of CD3 and CD4 T cells, and (C) CD8 T cells by expression of CD3 and CD8 T cells.

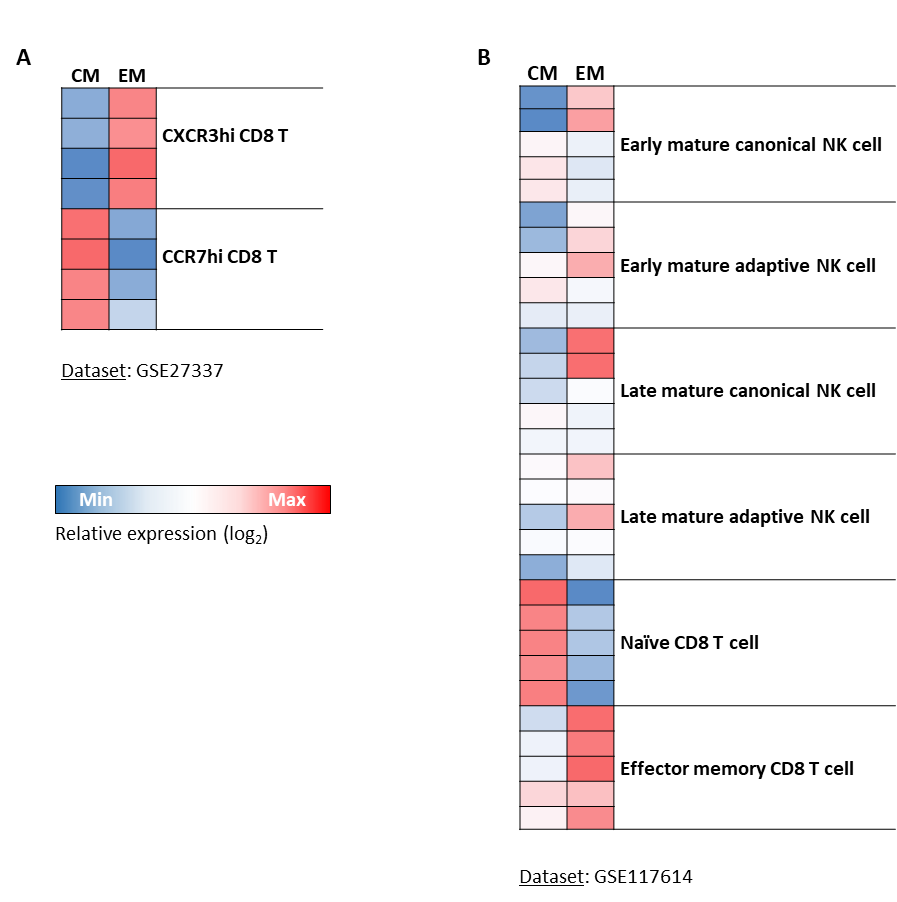

Dataset: GSE27337

Relative expression (log_2_)

**Min	Max**

**Supplementary Figure 5: Assessment of sensitivity and specificity of central memory (CM) and effector (EM) CD8 T cell modules.** The expression of CM and EM CD8 T-cell transcriptional modules in (A) CXRC3hi or CCR7hi CD8 T cells (GEO Omnibus dataset GSE27337) and (B) sorted NK cell and CD8 T cells (GEO Omnibus dataset GSE117614).
**A**

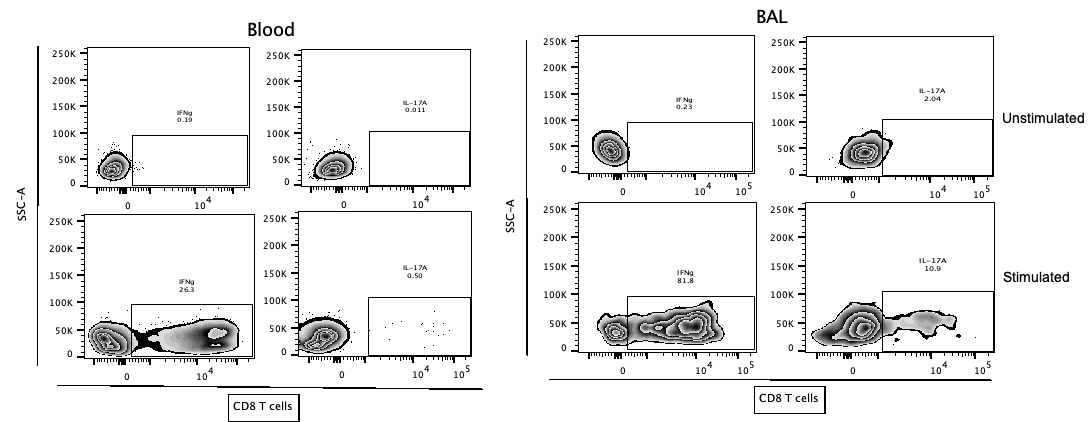

**B	C**

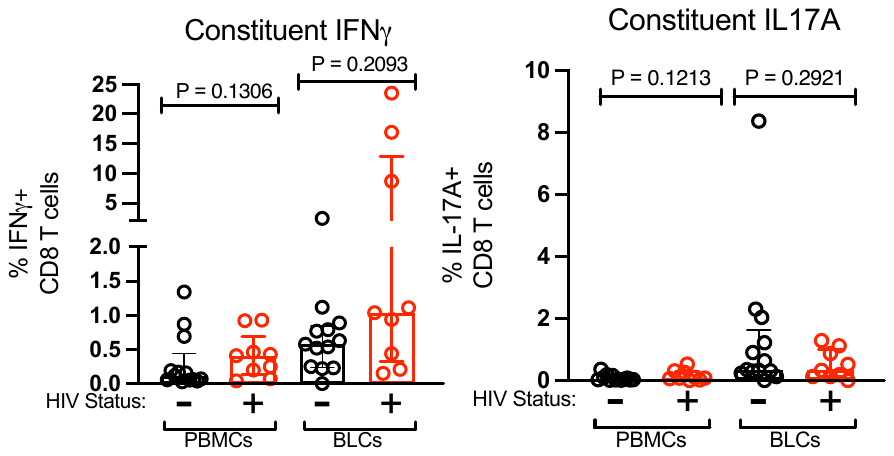

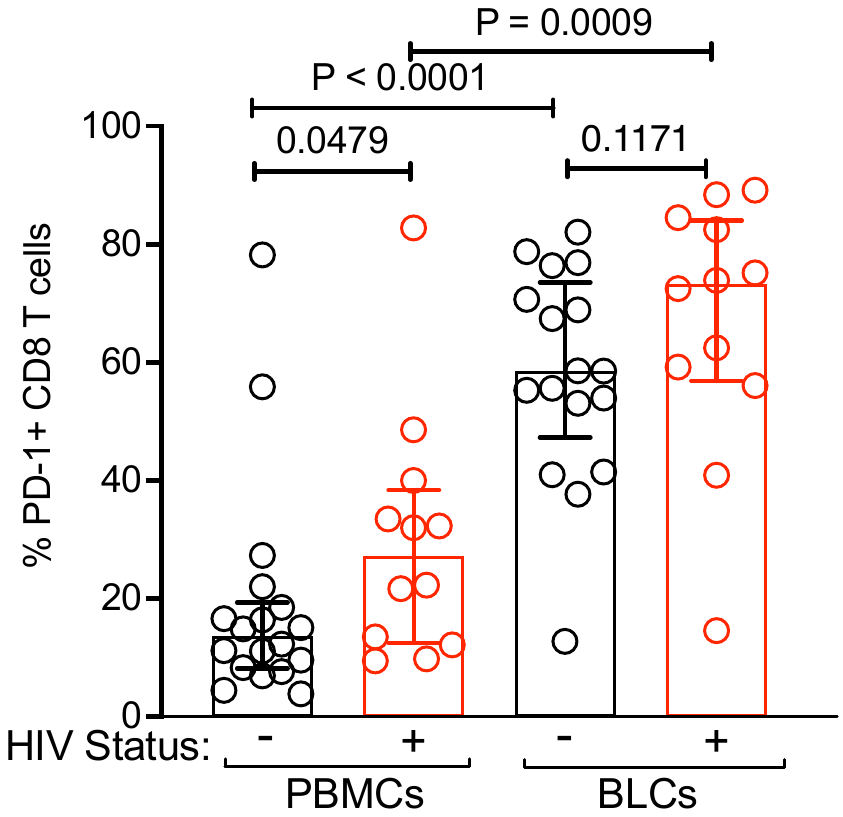

**Supplementary Figure 6: Blood and BAL samples IFN-γ** **and IL-17A expression in HIV infection. A.** Gating strategy showing a representative plot of the CD3+ and CD8+ live T cells, and their proportion of cells producing IFN-γ and IL-17A in either blood or BAL samples and under-stimulated (PMA/Ionomycin) and nonstimulated condition. B. Intracellular cytokine staining showing the percentage of CD8+ T-cells that produce IFN-γ and IL-17A under no mitogen stimulation condition in samples from blood and BAL compartment in PLWH and those without HIV. C. Protein level expression of PD-1+ CD8+ T-cells from blood and BAL samples in PLWH and those without HIV.
