## Supplementary Methods for "Dysfunctional effector memory CD8 T cells in the bronchoalveolar compartment of people living with HIV"

**Study subjects and samples acquisition procedures**

Bronchoalveolar lavage (BAL) fluid and paired peripheral blood samples were sourced from a research bronchoscopy study called Phefumula, based at Africa Health Research Institute (AHRI) in Durban, South Africa. This study was approved by the University of KwaZulu-Natal Biomedical Research Ethics Committee (BREC; reference numbers BF503/15 and BE037/12) and the Partners Institutional Review Board.

This cohort has been described by Muema et al, 2020 (1) and Khuzwayo et al, 2021 (2). Briefly, participants were recruited and consented to screen tests to assess eligibility for the research bronchoscopy study. Exclusion criteria for this study included a prior diagnosis of TB disease or any other co-morbid disease, smoking or pregnancy. People who were known to be HIV negative or who had newly diagnosed, and previously untreated HIV infection were eligible for screening. People with newly diagnosed HIV were referred for immediate antiretroviral therapy according to South African Department of Health guidelines. During the screening process, participants were screened for latent TB status using QuantiFERON-TB Gold Plus (QFT-Plus, Qiagen), HIV status was evaluated using 4^th^ generation HIV antibody/antigen Enzyme Linked-Immunosorbent Assay (ELISA) testing, HIV RNA quantitative viral load and CD4 T-cell count were also quantified. A chest X-ray and sputum GeneXpert were also performed to exclude active TB cases. People who were either HIV-negative or HIV-positive and ART-naïve, had a positive QFT result, and whose blood test met safety criteria for research bronchoscopy (hemoglobin > 10 g/dL, platelet count > 150 and international normalized ratio (INR) > 1) were offered participation in the research bronchoscopy study. On the day of the bronchoscopy, a peripheral blood draw was conducted. The research bronchoscopy procedure has been described previously (3); briefly after administration of topical lidocaine to the vocal cords and midazolam, the pulmonologist wedged the bronchoscope in the right middle lobe, instilled 200 mL of normal saline and collected lavage fluid for processing.

**Sample processing**

The paired blood and BAL samples were concurrently collected and subsequently transported to the research laboratory in less than 2 hours for processing. PAXGene tubes were immediately frozen. A portion of whole blood and BAL samples were assessed for differential cell count using histochemistry and compound microscopy to count for alveolar macrophages/monocytes, eosinophils, lymphocytes and neutrophils per sample type (Supplementary Figure 1). Peripheral blood mononuclear cells (PBMCs) were isolated from the bloods using the standard Histopaque^®^ (Sigma-Aldrich) gradient centrifugation method. A 1.5 mL portion of the whole BAL samples was centrifuged (1500 g x 10 mins), and the pellet was resuspended in 100 uL RNALater stabilizing reagent (Sigma-Adrich) and frozen. The remaining BAL sample was then filtered through a 40 µm filter, centrifuged (1500 g x 10 mins) and the cell pellet resuspended in 10% of the sample in saline buffer volume with completed RPMI media (RPMI media supplemented with 5% fetal bovine serum, 1% penicillin/streptomycin, 1% HEPES buffer and 1% amphotericin). When sufficient live mononuclear cells were available from blood and BAL samples, they were subjected to (Phorbol 12-myristate 13-acetate (PMA) (25 ng/mL)/Ionomycin (500 ng/mL)) stimulation for 6 hours at 37^o^C in 96-well microplates and monoclonal antibody staining (Supplementary Table 1).

**Preparation of RNA-seq libraries and sequencing**

Whole compartment: Ten pairs of blood and BAL samples from HIV-negative people and 10 pairs of blood and BAL samples from HIV-positive people (Supplementary Figure 1) were used to prepare RNA-Seq libraries. Preserved blood and BAL samples were thawed from -80^o^C storage to room temperature (RT). RNA was extracted from the Blood PAXgene® tubes using PaXgene Blood RNA kit (Qiagen) according to the manufacturer's recommendations. Briefly, PAXgene blood samples rested at RT for 2hrs to facilitate cell lysis and then centrifuged and the pellet was washed. Proteinase K was used to remove proteins from the samples and the sample passed through a Shredder spin column and DNAse I treated to homogenize and remove any genomic DNA. Filtrate was subsequently transferred into the RNA Spin column to isolate the total RNA in each sample. RNA extraction from the BAL samples was done using RNAEasy Micro Kit (Qiagen) following manufacturers protocol. Briefly, the BAL cells in RNALater reagent were pelleted and resuspended in 1% β-mercaptoethanol RLT lysis buffer (Qiagen) and RNA extraction and continued with extraction as above. The concentration and integrity of the extracted RNA was assessed using NanoDrop Lite Spectrophotometer (Fisher Scientific) and 4200 TapeStation System (Agilent Technologies), respectively. RNA-Seq libraries were prepared as described by Muema et al, 2020 (1). Briefly, RNA from both blood and BAL samples was enriched for messenger RNA using the NEBNext^®^ Poly(A) mRNA Magnetic Isolation Module (New England Biolabs). Libraries were prepared using NEBNext^®^ Ultra RNA Library Prep Kit for Illumina (New England Biolabs) and barcoded using NEBNext Multiplex Oligos for Illumina kit (New England Biolabs). Library quality was assessed using the DNA TapeStation (Agilent). RNA library concentrations were determined using a Qubit dsDNA assay kit (Thermo Fisher). A pool of barcoded libraries with equal molarities was then sequenced on an Illumina NextSeq 500 platform targeting 10 million reads per sample. After an initial sequencing run that included all samples, selected samples were re-sequenced to achieve goal sequencing depth.

Sorted cell populations: In participants with sufficient live cells from BAL compartments, adherent and non-adherent cells were separated by plastic adherence on 6 well plates (CLS3335) at 37^o^C for 1hr. Non-adherent cells were stained using Live/Dead™ Fixable Aqua (Life Technologies, L34957), CD3-BV650 (BioLegend, 317324), CD4-BV711 (BioLegend, 317440), CD14-APC-Cy7 (BioLegend, 325620), PD-1-BV421 (BioLegend, 562516 ), TIM-3-BV785 (BioLegend, 345031) and CD8-PE Texas Red (Invitrogen, MHCD0817). Adherent cells were scrapped from the plastic. Both samples were acquired on a BD FACSAria III (BD Biosciences) cell sorter to isolate CD4 T cells (defined as single, live, lymphocytes, CD3+, CD4+), CD8 T cells (defined as single, live, lymphocytes, CD3+, CD8+), MAIT cells (defined as single, live, lymphocytes, CD3+, CD4-, MR1-5OP-RU tetramer+), monocytes (defined as single, live, lymphocytes, CD3-, CD14+) and B-cells (defined as single, live, lymphocytes, CD3-, CD14-, CD19+) and alveolar macrophages (sourced from unstained adherent cells, defined by size and complexity). The gating and sorting (done for some BAL samples from PLWH) strategy is shown in Supplementary Figure 2. The purity of sorted cells was confirmed to be >95% for a test sample. Using single-cell sorting mode, ‘mini-populations’ of 100 cells were collected into 50 µL of RLT buffer (Qiagen) with 1% β-mercaptoethanol and subsequently stored at -80^o^C. Subsequent RNA isolation and library preparation from these samples was conducted using Smart-Seq and Smart-SeqII approaches as described in Trombetta et al, 2014 (4). Briefly, RNA from lysed mini-populations were isolated using SPRI paramagnetic bead technology (RNAClean XP, Beckman Coulter), and cDNA was prepared by targeting RNA with poly-A tail ensuring reverse transcription of mRNA using Maxima First Strand cDNA Synthesis Kit (Thermo Fisher Scientific, K1641). Whole transcriptome amplification was performed and cleaned using AMPure XP beads (Beckman Coulter). RNA libraries were then prepared using Nextera^®^ XT library preparation kit (Illumina).

Library quality was assessed using the DNA TapeStation (Agilent). RNA library concentrations were determined using a Qubit dsDNA assay kit (Thermo Fisher). A pool of barcoded libraries with equal molarities was sequenced on an Illumina NextSeq 500 platform. After an initial sequencing run that included all samples, selected samples were re-sequenced to achieve goal sequencing depth.

**Transcriptomic data extraction and assessment**

The RNASeqPipelineR package (https://github.com/RGLab/RNASeqPipelineR) was used to perform alignment, quantification and annotation of the RNA sequencing data (RNA-seq) from the whole and mini-population RNA-Seq libraries. The RNA-seq data were aligned against the human genome (hg38) using STAR (v2.4.2a) (5), and gene expression quantification was performed using RSEM (v1.2.22) (6). Genes with less than 15 nonzero read counts were discarded, leaving 19,720 expressed genes for the analysis. Libraries (samples) with less than 500,000 reads; 10,000 detected genes; an alignment rate < 75% and an exon rate < 50%; and clear sample annotation were confirmed using NGSCheckMate.

Differential gene expression of RNA-Seq data was analysed with DESeq2 and SARTools packages, using false discovery rate (FDR) <0.05 (7). Pathway analysis was performed using the Reactome database via InnateDB (8) (significantly enriched genes or pathways we defined by fold change > 1 or < -1 and FDR q value < 0.05 (P value adjusted using Benjamini-Hochberg correction). Ingenuity pathway analysis (Qiagen) was used to predict upstream regulators of differentially expressed genes, with significant regulators determined by corrected P values < 0.05.

**Transcriptional module analyses**

The expression of transcriptional modules in RNAseq datasets was performed as previously described by Pollara et al, 2017, with data log2 transformed and presented as counts per million (9). Module expression scores were derived from the geometric mean expression of all constituent genes within a module. For CD4 T cells, CD8 T cells, mucosal-associated invariant T cells (MAITs) and NK cell modules, we utilized those derived from the bulk transcriptome of purified immune cells (10) or blood samples, which possess the greatest sensitivity and specificity for cognate purified cell types (9, 11) and have been validated to reflect relative immune cell frequency in vivo (9, 12). Transcriptional modules representing immune cells in BAL were derived from differentially expressed genes between sorted cell populations using DESeq2. For each cell type of interest, module constituent genes were derived from genes with significantly elevated expression in the cognate cell relative to specific other cell types (Supplementary Table 2). The CD4 were compared to CD8 T cells and MAIT cells, while the CD8 T cells were compared to CD4 T and MAIT cells and the rest of the prepared modules were defined by comparing cognate cell type relative to all others (e.g. C14+ monocytes compared to CD4, CD8 T-cells, MAITs and alveolar macrophages). The sensitivity and specificity of these modules were internally validated from the BAL transcriptome from which they were derived (Supplementary Figure 3A) and externally validated using an independent dataset of sorted immune cell types (ArrayExpress dataset E-MTAB-7143) (Supplementary Figure 3B).

Transcriptional modules reflecting the biology of central and effector memory (CM and EM) CD8 T cells were derived from CD8 T cells purified on the basis of CCR7 and CXCR3 expression, with CCR7^hi^CXCR3^lo^ representing CM and CCR7^lo^CXCR3^hi^ EM CD8 T cells (13). We focused on genes that were differentially expressed between CM and EM populations (2-sided t-test, P < 0.01 uncorrected for multiple testing) and >10-fold expressed between the 2 groups. Genes that fulfilled these criteria were used to compose CD8 CM and EM transcriptional modules. Their specificity was initially determined by internal validation of expression scores on the dataset of module origin (GSE27337) and then using an independent external dataset (GSE117164) of purified CD8 T cell subsets and NK cells (14). These revealed high sensitivity and specificity of the CM and EM transcriptional modules for their cognate cell types (Supplementary Figure 5).

**Transcriptional linear regression analyses**

To adjust for the contribution of different cell type frequencies on differential gene expression in the BAL transcriptome, a linear regression was performed for each gene of the expression matrix using each module expression score to generate a vector of cell counts. The slope and intercept of this vector were used to predict the expression of all genes, and thus calculate residuals for each gene in the matrix for all samples. The expression matrix of residuals generated was then used for differential gene expression in the same way.

**Flow cytometry**

Mononuclear cells from both blood and BAL samples were counted and viability was assessed and confirmed to be >90% using trypan blue (Sigma-Aldrich). To assess cell lineage distribution, phenotypes, and functionality of immune cells in blood and BAL compartments with HIV infection we stained mononuclear cells using 2 panels of fluorescently labeled monoclonal antibodies. Panel 1 (phenotypic panel): Same as the sorting panel above. Panel 2 (intracellular cytokine staining panel): Live/dead-amcyan (Life Technologies), CD3-Alexa700 (BioLegend, 300424), CD4-BV711 (BioLegend, 317440), CD8-APC-Cy7 (BioLegend, 344714), granzyme b-Alexa647 (BioLegend, 515406), IFN-γ-PE/Dazzle 549 (BioLegend). Staining using Panel 2 was done after PMA/Ionomycin (25/500ng/mL) stimulation. The BDFACSAria III (BD Biosciences) flow cytometer was used to acquire samples. Compensation and rainbow beads were used as standards to minimize day-to-day variability. FlowJo v10.8.1 (FlowJo, LLC) was used for flow cytometry analysis. Comparisons in the flow cytometry data between HIV-negative and HIV-positive people were evaluated using Mann-Whitney U test.

**Methods References:**
